# AltraFlowSOM: A Semi-Supervised Framework for Imaging Mass Cytometry Phenotyping

**DOI:** 10.64898/2026.08.31.748226

**Authors:** Akhiya Anilkumar Rekha, Eleonore Bettacchioli, Christelle Le Dantec, Patrice Hemon, Pierre Emmanuel Jouve, Sophie Hillion

## Abstract

Imaging Mass Cytometry (IMC) enables the simultaneous quantification of 40+ protein markers at single cell resolution in tissue, however biologically faithful phenotyping at scale remains a critical bottleneck. Unsupervised clustering fragments coherent populations or conversely merges biologically incoherent ones into a single cluster, supervised classifiers impose a closed vocabulary, and the presence of rare subsets (encoding clinically relevant biology) in conjunction with abundant subsets may be detrimental to detection performances. We present AltraFlowSOM, a semi-supervised extension of FlowSOM that embeds partial expert annotations directly into self-organizing map training via a two-layer SuperSOM architecture, balancing label-guided topology anchoring with unsupervised discovery. By anchoring the map to biologically labelled reference points, AltraFlowSOM circumvents the canonical dependency between batch correction and clustering. Evaluated under Leave-one-out cross validation on two independent IMC cohorts, Lupus Nephritis (n=22 ROIs) and Sjögren syndrome (n=10 ROIs), AltraFlowSOM outperformed all unsupervised and supervised baseline on Adjusted Rand Index, F1 scores (macro and weighted), weighted purity and in the identification of rare populations. The median Treg cell recovery exceeded that of all comparator methods. AltraFlowSOM resolves the scalability-alignment-discovery trilemma, by establishing a semi-supervised SOM as a generalizable method for high dimensional IMC phenotyping.

## 1. Introduction

High-plex spatial proteomics techniques, such as MIBI (Multiplexed Ion Beam Imaging), CODEX (Co-detection by indexing), cycIF (Cyclic Immunofluorescence), and Imaging Mass Cytometry (IMC) generate high-dimensional spatial data. Among these methods, IMC uses metal tagged antibodies detected by time-of-flight cytometry to overcome the multiplexing limitations of traditional immunohistochemistry and immunofluorescence enabling the simultaneous staining, acquisition and analysis of 40+ markers per tissue section. This allows for the comprehensive characterization of cellular composition, cell-state programs, and spatial organization across a broad range of pathological contexts, including tumor microenvironments, autoimmune diseases, and transplant rejection (Louis Sam Titus et al. 2023).

Despite this resolution, IMC analysis remains computationally challenging primarily because of tissue complexity and technical variability (Giesen et al. 2014). Marker panels are heterogeneous in information content : lineage markers dominate distance metrics, while functional state markers carry subtler signals leading to under-resolution of activation or effector programs (Nowicka et al. 2017, Liu et al. 2019). Batch effects from staining intensity, instrument drift, and sample handling can shift marker distributions (Tan et al. 2020), and segmentation artifacts shared across multiplexed imaging platforms can distort marker profiles and obscure rare populations. These technical factors compound genuine biological heterogeneity, making it essential to distinguish between technical variation and true biological signals (Hunter et al. 2024).

Existing strategies have distinct and complementary limitations. Manual gating is interpretable and aligns closely with immunological reasoning (Qiu et al. 2011), yet it is subjective, time-consuming, and difficult to reproduce at scale. Unsupervised methods such as FlowSOM (Van Gassen et al. 2015) and PhenoGraph (Levine et al. 2015) are scalable and discovery-oriented, but are sensitive to preprocessing choices and prone to biological fragmentation, splitting coherent cell populations into multiple clusters. Supervised classifiers (e.g., Random Forest) (Spies et al. 2025) agree well with the curated labels, but cannot represent unknown or transitional states. Deep-learning approaches such as CellSighter (Amitay et al. 2023) and MAPS (Shaban et al. 2024) achieve expert-level accuracy but require fully labelled, panel-specific training data. Knowledge-based tools such as Tribus (Kang et al. 2025) avoid manual labelling but apply scoring post-hoc, precluding the discovery of unanticipated phenotypes. A cross-cutting limitation is class imbalance: rare but clinically critical subsets such as tissue-infiltrating regulatory, effector subsets, and disease-specific populations encoding disease activity and treatment response (Hartmann and Bendall, 2020) are systematically absorbed or fragmented regardless of the analytical paradigm.

To address this gap, we developed AltraFlowSOM, a semi-supervised extension of FlowSOM that integrates partial expert annotations directly into self-organising map training via a two-layer SuperSOM architecture. Evaluated on two independent IMC cohorts under leave-one-out cross-validation, AltraFlowSOM outperformed all unsupervised and supervised baselines in terms of Adjusted Rand Index (ARI), Macro-F1, and rare-population recall.

## 2. Materials and Methods

### 2.1 Datasets

#### 2.1.1 IMC cohorts

Two independent IMC cohorts comprising biologically distinct tissues were used to assess robustness across heterogeneous settings. Cohort 1 comprised AFA-fixed kidney sections (3 µm) from eight patients with lupus nephritis (LN). Twenty-four ROIs were acquired, of which two were excluded due to high background noise, yielding 22 ROIs for analysis using 34 markers. Cohort 2 comprised minor salivary gland biopsies (5 µm) from seven patients with primary Sjögren’s syndrome (SjD), with 10 ROIs (1 × 1 mm2), selected for analysis using 27 markers. The full clinical characteristics and marker information are provided in **Supplementary Tables S1 (LN) and S2 (SjD)**. All samples were acquired using Hyperion Imaging System (Standard BioTool).

#### 2.1.2 Cell Segmentation and Feature extraction

The IMC images were segmented to define single-cell boundaries prior to feature extraction. LN sections were segmented using QuPath (Bankhead et al. 2017) and SjD sections using YOUPI (Scuiller et al. 2023) leading to the identification of **131528** and **115523** cells respectively. IMC data were then processed following established preprocessing procedures (Windhager et al. 2023). Per cell marker intensities were extracted to form a cell by marker (expression matrix) matrix X∈ R^nxd^, where n is the number of cells and d is the number of markers. Background channels, segmentation artifacts and low quality channels were removed prior to downstream analysis. The complete workflow is illustrated in **Fig. 1**.

**Fig. 1.**
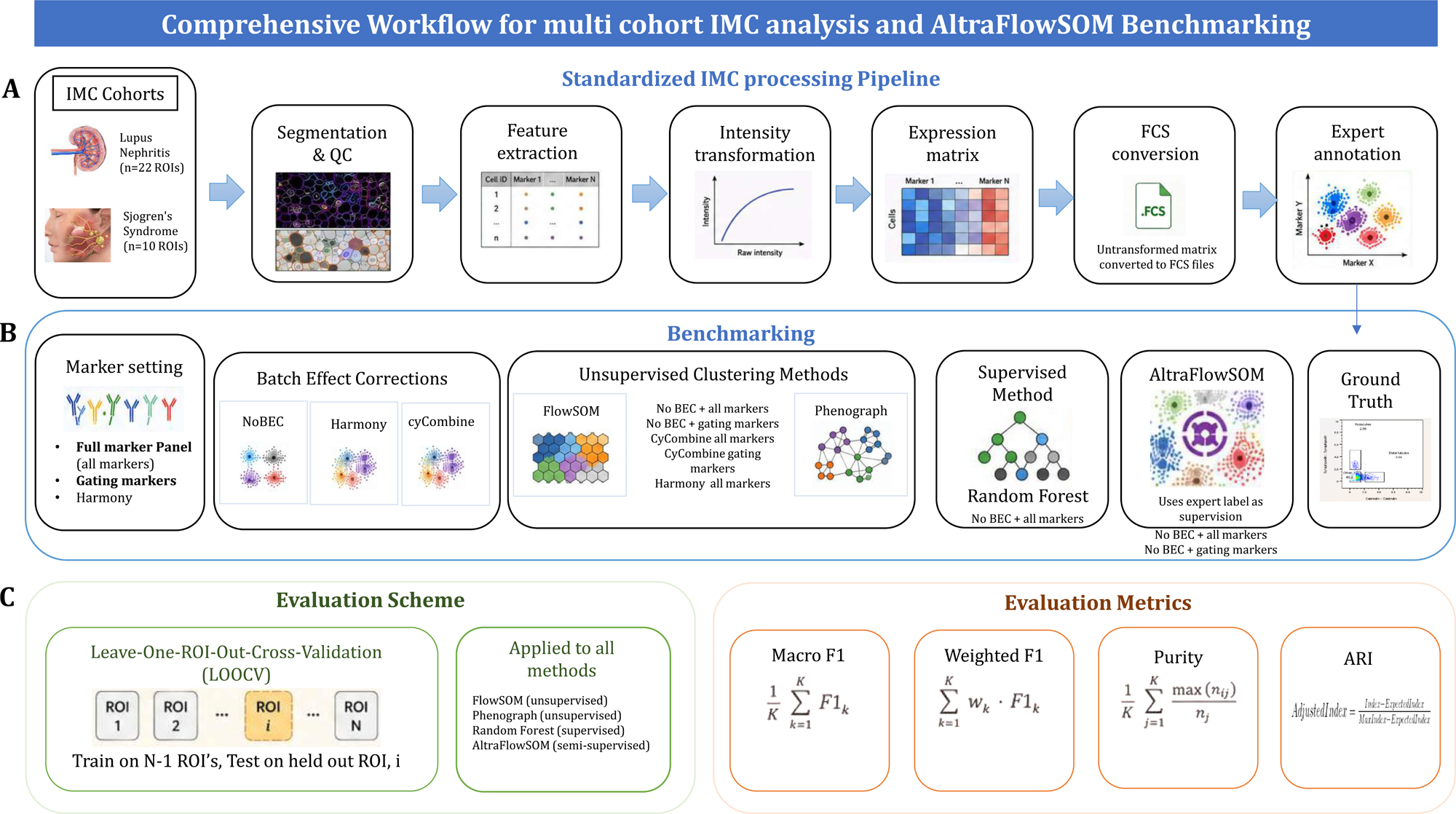
Comprehensive workflow for multi-cohort IMC analysis and AltraFlowSOM benchmarking. (A) Two IMC cohorts : Lupus Nephritis (n = 22 ROIs) and Sjögren’s syndrome (n = 10 ROIs), were processed through a standardized pipeline covering cell segmentation, single-cell feature extraction, and intensity transformation to generate cell-by-marker matrices. (B) Using both a full marker panel and a gating-marker subset, three batch-effect conditions were evaluated (NoBEC, Harmony and cyCombine) followed by unsupervised clustering with FlowSOM or Phenograph. In parallel, the intensity matrices were converted to FCS format and manually annotated in FlowJo to generate expert reference labels, which served as the supervision layer for AltraFlowSOM and as the ground truth for all benchmarking metrics. (C) All methods were evaluated using leave-one-ROI-out cross-validation (LOOCV); A Random Forest classifier was included as a supervised baseline. The performance was assessed using the Macro-F1, Weighted F1, mean purity, and ARI.

#### 2.1.3 Intensity transformation

Arcsinh cofactors spanning values 0.01 - 3 and logarithmic variants (log(1+x), log(2+x)) were systematically evaluated. A cofactor of 0.1 was selected because it yielded the best balance between reducing distributional skewness and preserving biological variation across both cohorts (Hunter et al. 2024). The transformed matrix was used directly for all downstream analyses.

#### 2.1.4 Batch Effect Correction

Batch-effect correction was treated as a primary experimental variable because inter-sample technical variation is a known source of performance differences in multi-patient IMC studies. Three conditions were evaluated for the transformed marker matrix:

- No correction (NoBEC)
- Harmony (Korsunsky et al. 2019) - PCA-based integration widely adopted in single-cell workflows. This was applied to the transformed matrix prior to clustering.
- cyCombine (Pedersen et al. 2022) which performs cluster-based integration and rank-informed normalization was designed for high-dimensional cytometry data. This was applied as a complementary condition to assess the robustness of the conclusions regarding the correction choice.

#### 2.1.5 Marker configurations

Two marker configurations were evaluated under all benchmarking conditions: **Marker Set A (all markers)** and **Marker Set B (gating markers)**, the latter corresponding to the marker subsets used by expert annotators during manual gating. Gating markers were defined a priori as those necessary to resolve the canonical immune and structural lineages present in each tissue. The complete panel and biological rationale for their selection are presented in **Supplementary Tables S1 and S2.**

#### 2.1.6 Manual gating reference labels

Manual gating labels constituted the ground-truth reference for all benchmarking metrics and the supervision layer for AltraFlowSOM. Prior to gating, the preprocessed single-cell expression matrix (untransformed) was exported to the FCS format to enable visualization and hierarchical gating in standard flow cytometry software. Gating was then performed in FlowJo® using a hierarchical two-dimensional gating strategy applied to Marker Set B, and reviewed independently by two expert annotators. Cells outside the canonical gate were assigned to an undefined category. These labels served as phenotype references for metric computation and partial labelling input for semi-supervised training. The complete gating hierarchy and gating strategy and representative gate plots for each annotated population are provided in the Results section **(Section 3.2, Fig. 4A and 4B)**.

### 2.2 AltraFlowSOM multi-layer semi-supervised clustering framework

We developed AltraFlowSOM, a semi-supervised (Robinson et al. 2026, Braga and Bassani 2018) clustering framework designed to integrate high-dimensional cytometry data with auxiliary biological priors. Its global workflow is strategically adapted from FlowSOM, a widely adopted framework for cytometry data analysis (Van Gassen et al. 2015). Following this established algorithm, AltraFlowSOM utilizes a 3-step hierarchical architecture (**Fig. 2**).

a. Topological Mapping and Training, based on the foundational Self-Organizing Map (SOM) theory developed by Kohonen, to condense high-dimensional cellular data into representative prototypes; ie; unlike standard FlowSOM, implementations which train a single SOM on marker expression layer alone, AltraFlowSOM employs a SuperSOM architecture that jointly trains on multiple parallel data layers including marker expression and label information, allowing topological organisation to be informed by both phenotypic and annotation structure simultaneously.
b. Metaclustering: Prototypes are subsequently grouped into higher-level clusters representing distinct biological populations
c. Prediction: Each individual cell is associated with its closest prototype and corresponding meta-cluster, enabling the scalable annotation of large cytometry datasets.

**Fig. 2:**
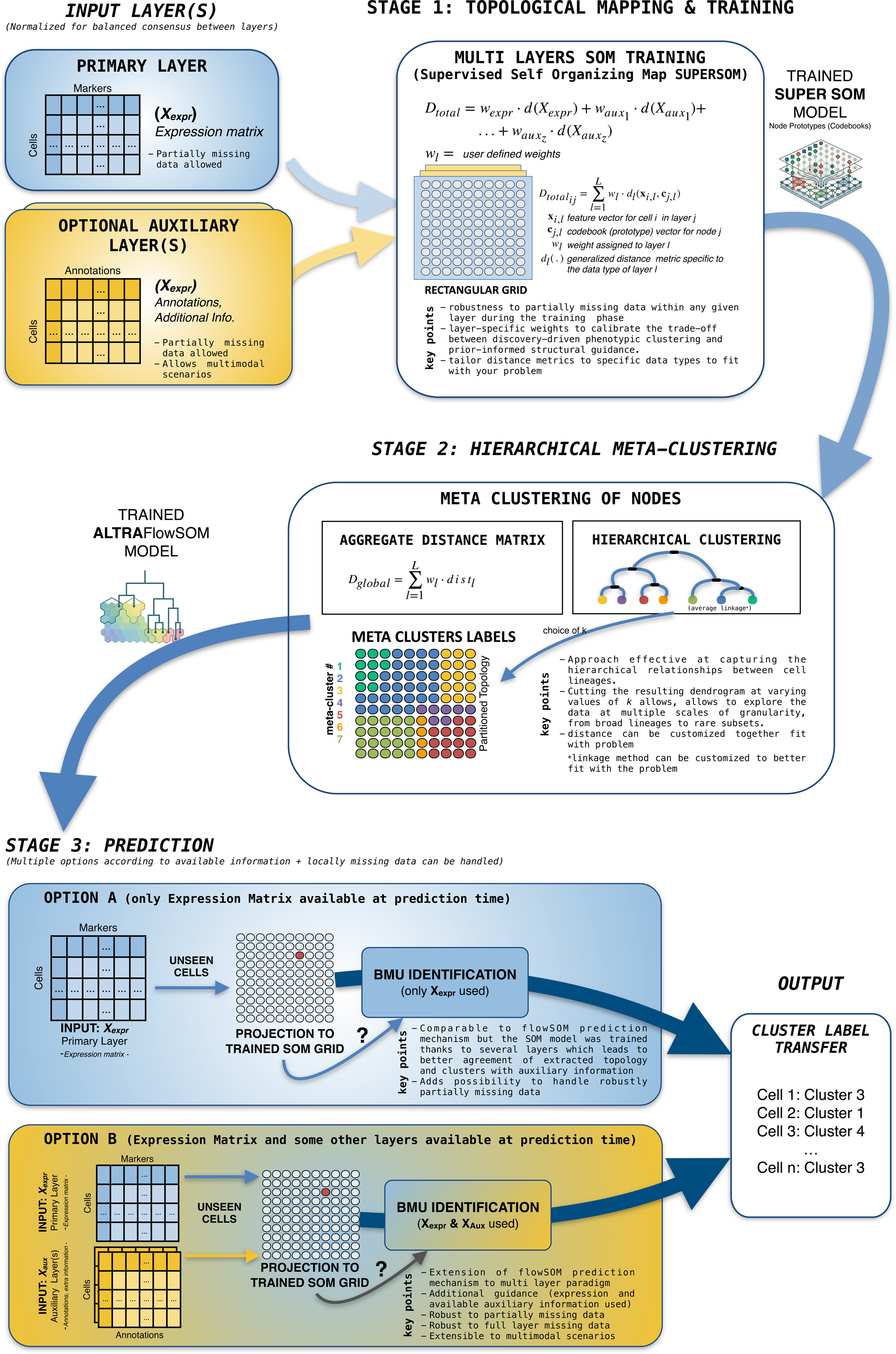
AltraFlowSOM framework description

The primary innovation of AltraFlowSOM lies in its transition from a single-layer unsupervised model to a multi-layer semi-supervised architecture. This is implemented through the Super-SOM (supersom) algorithm, an extension of the Kohonen map that allows for the simultaneous training of multiple heterogeneous data sources (layers) while maintaining a shared map topology (Wehrens and Kruisselbrink 2018, Wehrens and Buydens 2007) (The supersom algorithm is available as part of the kohonen R package, which can be accessed on CRAN at: https://cran.r-project.org/package=kohonen) .

The AltraFlowSOM’s architecture enables cytometry measurements to be jointly modeled with auxiliary biological priors, thereby guiding cluster identification using both data-driven and prior-informed information. The following sections detail each stage of the workflow.

#### 2.2.1 Topological mapping and training : Multi-layer training with robust handling of missing data

During the training phase, the cells are mapped onto a rectangular SOM grid. Unlike conventional SOM approaches, where the topology is learned exclusively from marker expression profiles, AltraFlowSOM enables the map to be jointly guided by multiple data layers. The algorithm identifies the Best Matching Unit (BMU) for each cell by minimizing a composite joint objective function that integrates information across all layers. For a given cell i and a node j, the combined distance D_total(ij)_ is defined as the weighted sum of the layer-specific distances:

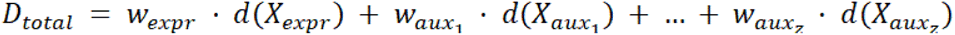

A critical feature of the supersom implementation is its native robustness to partially missing data within any given layer during the learning phase (Wehrens and Kruisselbrink (2018)Wehrens and Kruisselbrink 2018). When a cell presents incomplete entries, either in cytometry expression data or auxiliary data, the distance computation is dynamically restricted to the available features. This ensures that the model leverages all available data points without requiring biased imputation strategies. Normalization across layers ensures a balanced consensus between the intrinsic phenotypic distribution and the prior biological structure.

Therefore, the AltraFlowSOM multi-layer training architecture offers a modular and highly customizable approach. For instance, by adjusting layer-specific weights (w_l_), researchers can precisely calibrate the trade-off between discovery-driven phenotypic clustering and prior-informed structural guidance. Furthermore, the ability to tailor distance metrics (d_l_) to specific data types, combined with inherent robustness to partially missing information, ensures that the framework remains statistically sound across diverse experimental designs and incomplete datasets. The algorithm implemented in this study used two layers (expression data and annotation data) but this can be extended to three layers and beyond (while those layers might embed totally different data types their combined use is rendered possible). This flexibility allows AltraFlowSOM to serve as a bridge between purely unsupervised exploration and highly specific, biologically grounded classifications.

#### 2.2.2 Hierarchical Meta clustering

To transform the high-resolution SOM grid into interpretable cell populations, we performed a secondary clustering of the node prototypes. Consistent with the FlowSOM methodology (Van Gassen et al. 2015), this meta-clustering step reduces noise and clarifies population boundaries.

AltraFlowSOM constructs a global inter-node distance matrix by aggregating the distances across all layers used during the training phase. Hierarchical clustering is subsequently applied with average linkage to this matrix. This approach is particularly effective in capturing the hierarchical relationships between cell lineages. By cutting the resulting dendrogram at varying values of k, researchers can explore the data at multiple scales of granularity, from broad lineages to rare subsets.

#### 2.2.3 Prediction with partially available information

AltraFlowSOM supports “asymmetric prediction” (prediction with all layers / prediction with missing layers = prediction based on a subset of layers; also note that for the 2 previous scenarios AltraFlowSOM can accommodate locally missing information for any layer taken into account), allowing for the classification of new, unseen data even when auxiliary layers are entirely or partially missing (Wehrens and Kruisselbrink 2018, Wehrens and Buydens 2007). This is particularly relevant when extensive metadata or expert annotations are available for a subset of the data but are absent in another subset. For this latter subset, AltraFlowSOM projects the cells onto the trained SOM using only the available layers (e.g., marker expression). Because the auxiliary information was used to “anchor” the node positions during training, the BMU identification in the restricted feature space still benefits from the semi-supervised guidance. The cell then inherits the meta-cluster label associated with its BMU, effectively transferring the biologically informed structure established in the training phase to the incomplete datasets.

### 2.3 Benchmarking baselines

AltraFlowSOM was benchmarked against three comparator methods covering the full spectrum from unsupervised to fully supervised phenotyping, evaluated under, preprocessing settings, marker configurations, batch-correction conditions, and the same leave one ROI out cross validation (LOOCV) scheme (**Section 2.4**).

#### 2.3.1 FlowSOM

FlowSOM (Van Gassen et al. 2015) served as the primary unsupervised comparator; we applied the standard FlowSOM workflow to the transformed expression matrix using only the expression layer X_expr_. After training the FlowSOM self-organizing map, the algorithm’s standard meta-clustering step was applied to the SOM codebook vectors using agglomerative hierarchical consensus clustering with average linkage. Meta-cluster solutions were then evaluated for k ={10, 20….90}. The test set cells were mapped to their nearest node by expression distance and the labels were propagated by majority vote.

#### 2.3.2 Phenograph

Phenograph (Levine et al. 2015) was included as an alternative unsupervised baseline using k-nearest-neighbour graph construction (k = 30) followed by Louvain community (Blondel et al 2008) detection. The resolution parameter was varied to span the same approximate cluster range as FlowSOM for comparison purposes.

#### 2.3.3 Random Forest

A (basic) Random Forest classifier (O’Brien and Ishwaran 2019, Cheng et al. 2023, Chen et al. 2004, Riese et al. 2020) was included as a fully supervised reference. Under LOOCV, the classifier was trained on all labelled cells from the n-1 training ROIs and applied to the held-out test ROI. As a supervised approach, Random Forest requires (a) all populations to be defined a priori and (b) a representative fully annotated training dataset to be available. Its performance under LOOCV is expected to degrade for rare populations because of population abundances across ROI-level folds and because its parameter settings were not tuned to handle class imbalance efficiently.

### 2.4 Evaluation protocol

#### 2.4.1 Cross validation strategy

ROI-level leave-one-out cross-validation (LOOCV) was used: at each fold, one ROI was held out as the test set and the remaining n-1 ROIs formed the training set, testing generalization across tissue-region heterogeneity. To mitigate stochastic variation from SOM initialization, five independent runs were performed per sample per clustering resolution and all reported metrics were medians or means across these runs.

#### 2.4.2 Performance metrics

Four complementary metrics were computed at each clustering resolution k and each marker condition, using manual gating labels as the reference standard:

1. The **Adjusted Rand Index** (ARI) measures the global pairwise agreement between the predicted and reference partitions, corrected for chance. An ARI of 1 indicates perfect agreement and an ARI of 0 indicates agreement no better than random.
2. **Weighted purity** was calculated as the cluster-size-weighted mean of these cluster-level purities, giving larger clusters proportionally greater influence on the final score. Weighted purity increases monotonically with cluster number; it must be interpreted alongside Macro-F1 or ARI to indicate within-cluster homogeneity.
3. **Macro F1** is the unweighted mean of the per-class F1 scores, treating each cell-type class equally regardless of its frequency. The macro F1 is insensitive to class imbalance and is therefore the primary metric for assessing the recovery of rare populations.
4. **Weighted F1** is the cluster size weighted mean of the per-class F1 scores, which reflects the overall recovery dominated by abundant populations.

#### 2.4.3 Rare functional population evaluation

The recovery of rare functional populations was evaluated separately from overall clustering performance, as aggregate metrics such as Macro F1 average performance across all classes and cannot isolate the recovery of any single population of interest. Because regulatory T cells (Tregs) are a biologically important but numerically sparse population in both cohorts, Treg-specific recall was recorded separately at each clustering resolution and marker condition.

## 3. Results

AltraFlowSOM was evaluated on the two independent IMC cohorts under ROI-level LOOCV (See Methods **Section 2**). For each held-out sample, five independent runs were performed at each clustering resolution within a range of 10-90 clusters. Across all held-out samples, this yielded 110 independent run-sample combinations for Cohort 1 (5 runs × 22 ROIs) and 50 for Cohort 2 (5 runs × 10 ROIs) per method configuration, with each combination evaluated across the full resolution range. AltraFlowSOM was benchmarked against FlowSOM (FS) and Phenograph (PG), each tested under three batch-effect correction (BEC) conditions (NoBEC, Harmony and cyCombine) and two marker configurations (gating markers and all markers). A supervised Random Forest (RF) classifier served as an upper-range reference.

### 3.1 AltraFlowSOM outperforms unsupervised and supervised baselines across two independent IMC cohorts

AltraFlowSOM achieved the highest ARI and Macro F1 across both cohorts, all clustering resolutions, and both marker configurations **(Fig. 3A and 3B; Suppl. Fig. 1A and 1B**), outperforming Random Forest despite RF’s despite RF’s supervised mode of action, an outcome anticipated by the class imbalance argument in **Section 2**. Random Forest, being a supervised approach, is only able to associate each instances with a unique pre-defined label and therefore yields a single fixed value per metric (ARI ∼0.32, Macro F1 ∼0.28, Weighted F1 ∼0.49, weighted purity ∼0.565), shown as a flat reference line across all panels rather than a curve over the 10-90 cluster range.

**Fig. 3:**
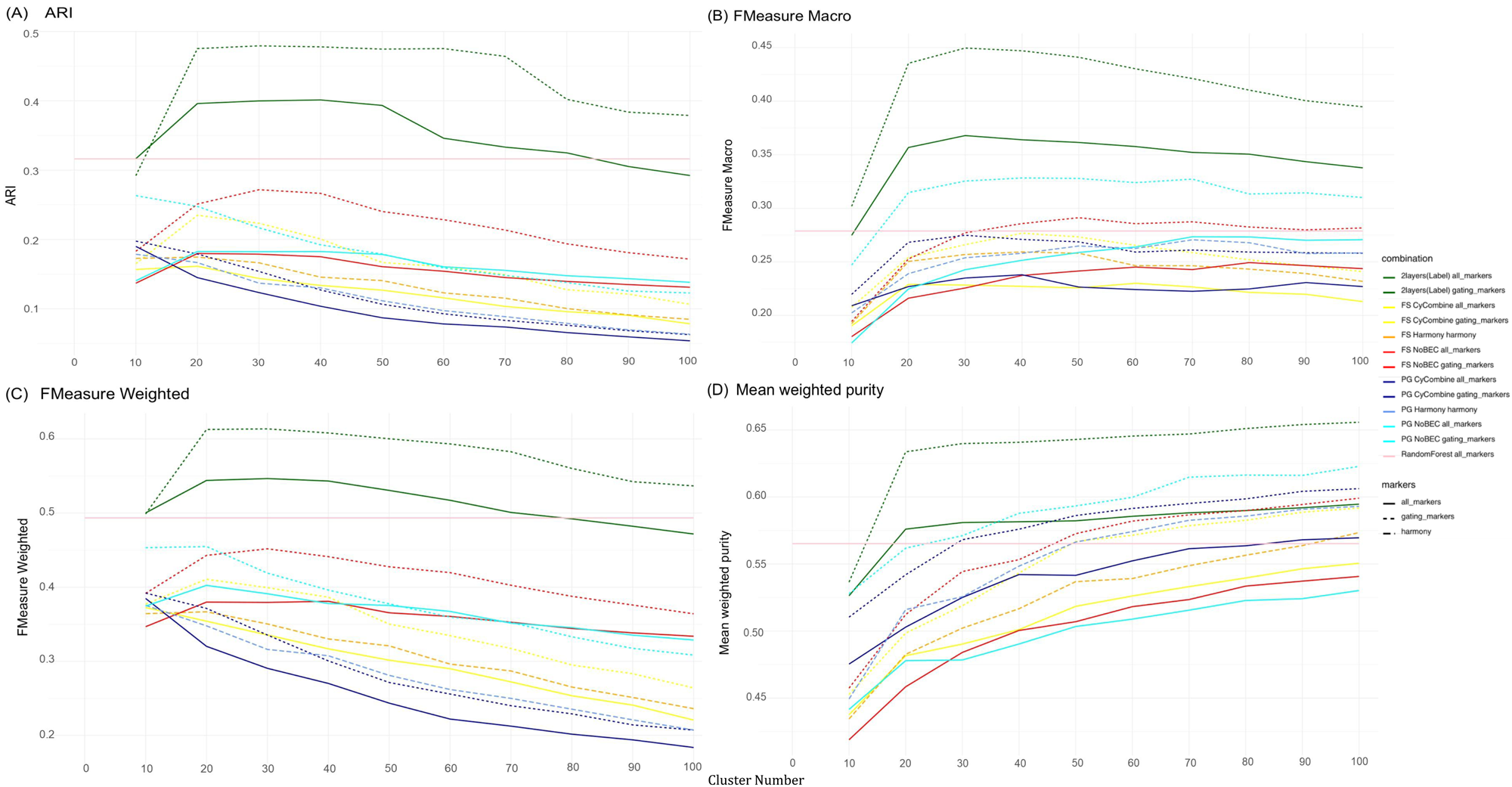
Line plots showing clustering accuracy across methods, marker configurations, and cluster resolutions. Panel (A) shows Adjusted Rand Index (ARI), panel (B) shows Macro-F1, panel (C) shows Weighted F1, and panel (D) shows mean purity for Cohort 1.

On ARI **(Fig. 3A)**, AltraFlowSOM (gating markers) peaks ∼0.48 at 20-40 clusters, with all-markers close behind (∼0.40); every other method (FlowSOM, Phenograph, Random Forest) stays within 0.05-0.27 across all resolutions and batch corrections, this clear, uniform margin reflecting better overall agreement between predicted and true cluster assignments. The same ranking holds for Macro F1 **(Fig. 3B)**: AltraFlowSOM (gating markers) peaks ∼0.45 at ∼30 clusters, the next-best method (PG NoBEC) only reaches ∼0.33, and the remaining methods sit at 0.22-0.29. Because Macro F1 weights each class equally regardless of size, this gap specifically reflects stronger recovery of rare populations, not just abundant ones. On Weighted F1 **(Fig. 3C)**, AltraFlowSOM still leads (gating ∼0.61, all-markers ∼0.54), but the competitor’s spread compresses to ∼0.40-0.45 (PG/FS NoBEC). Since Weighted F1 down-weights rare-class errors by frequency, this narrowing is expected to measure something closer to majority-class performance, where methods differ less. On weighted purity **(Fig. 3D)**, AltraFlowSOM (gating markers) leads throughout (∼0.53→0.65 max), but the all-markers variant is overtaken by PG NoBEC past ∼60 clusters, with several competitors converging near it by 90 clusters. Weighted purity rewards finer partitioning regardless of biological relevance, so any method’s purity score rises with more clusters; this convergence at high resolution is a known property of the metric. Across all four metrics, AltraFlowSOM consistently outperforms FlowSOM, Phenograph, and Random Forest, with the largest margins on ARI and Macro F1 the metrics most sensitive to correct, class-balanced cluster assignment.

### 3.2 Phenotype to Cluster agreement

To decompose the clustering performance at the population level, we first examined the distribution and absolute cell percentage of the annotated cell types in both Cohorts (**Fig. 4C and Suppl.Fig. 2C**). In cohort 1 (LN), the dataset was characterized by a pronounced class imbalance: with structural compartments (Other_7, Other_8, CD138+ Tubules and CD163+ Macrophages) numerically dominating (**Fig. 4A and 4B)**, whereas functionally important immune subsets such as Tregs represented less than 1% of the cells. A comparable imbalance was also observed in Cohort 2 (**Suppl. Fig. 2A and 2B**).

**Fig. 4:**
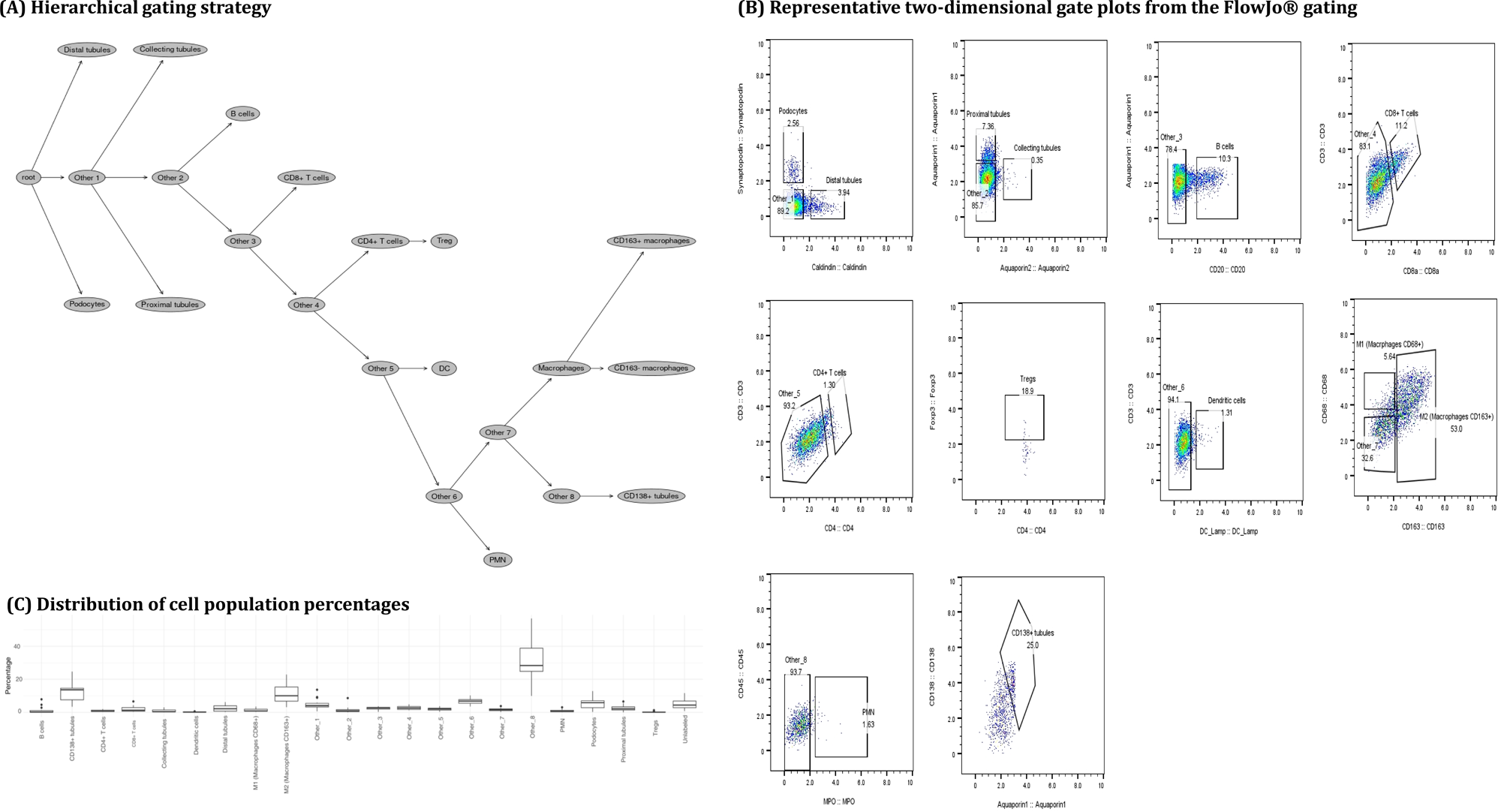
(A) Hierarchical gating strategy used for expert annotation of IMC single-cell data in FlowJo®. The gating strategy was applied on the untransformed expression matrix exported to FCS format. (B) Representative two-dimensional gate plots from the FlowJo® gating session, illustrating the marker combinations and gate boundaries used to define each annotated population. Gate labels correspond to the cell type nomenclature used throughout the benchmarking analysis. (C) Distribution of cell population percentages across 23 annotated immune and non-immune cell types in Cohort 1 (LN), illustrating the pronounced class imbalance that motivates the use of Macro-F1 as the primary evaluation metric. For the heatmaps shown below, the categories other_1, other_2.. were merged into a single other/Not defined category.

To investigate per-population performance, predicted cluster assignments were compared with manual gating labels using heatmaps at k = 30 with gating markers across both cohorts (**Fig. 5, Suppl. Fig. 3**). Two complementary normalizations were applied: row normalization (**Fig. 5A, Suppl. Fig. 3A**), giving each population’s recall (fraction of its cells assigned to each cluster), and column normalization (**Fig. 5B, Suppl. Fig. 3B**), giving each cluster’s purity (its composition by annotation). Given the class imbalance in **Fig. 4** and **Suppl. Fig. 2C**, both must be read jointly, as neither alone captures recall and purity simultaneously.

**Fig. 5:**
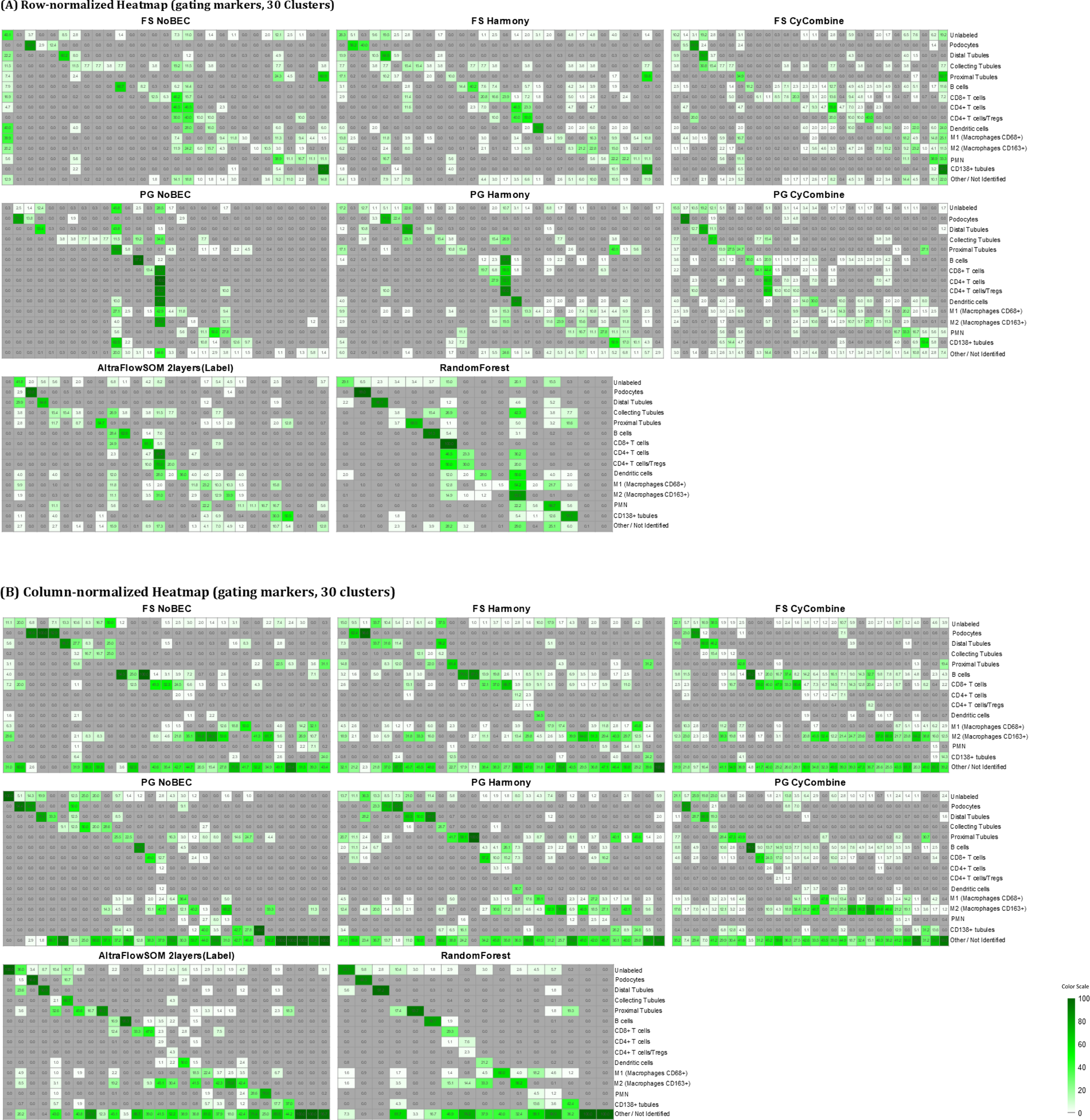
Agreement between AltraFlowSOM-predicted and manually annotated phenotypes at 30 clusters using gating markers in Cohort 1 (LN). (A) Row-normalised heatmap showing annotation recall. (B) Column-normalised heatmap showing cluster purity.

Each row reflects the fraction of a given manually annotated cell type that is captured by each cluster. For FlowSOM and Phenograph (across all batch-correction conditions), most rows show recall diffusely distributed across many clusters, with no single cluster capturing a majority of any population, indicating that these unsupervised methods fragment most ground-truth types across multiple clusters. Random Forest, as a directly trained supervised classifier, shows the sharpest recall concentration overall, with abundant and rare types alike routed predominantly into a single cluster. AltraFlowSOM achieves recall concentration approaching that of Random Forest for several populations, including Podocytes, Distal Tubules, Proximal Tubules, B cells, and CD8+ T cells, each mapping the majority of its ground-truth cells into one or two dedicated clusters, despite being trained without direct supervision on the full label set.

Each column reflects the cell-type composition of a single cluster. For FlowSOM and Phenograph, most clusters contain mixed contributions from multiple ground-truth types, with few columns reaching high single-type purity; rare populations (CD4+/Tregs, PMNs, dendritic cells) are essentially undetectable as pure clusters under either method. AltraFlowSOM produces several high-purity clusters corresponding to structural and lineage-defining populations, indicating that incorporating label-guided information helps AltraFlowSOM form dedicated, compositionally clean clusters for each cell type, rather than mixed populations. Random Forest again shows the highest purity overall, consistent with supervised, high-confidence class assignment.

### 3.3 Purity-Macro F1 trade-off

To summarize the homogeneity recovery trade-off across all method configurations at a single resolution, weighted purity and Macro-F1 were plotted jointly at k = 30 for both cohorts (**Fig. 6A and 6B**). AltraFlowSOM with gating markers occupied the upper-right quadrant (Macro-F1 0.42 and weighted purity of 0.66), demonstrating that the semi-supervised design resolves the trade-off that constrains all unsupervised methods.

**Fig. 6:**
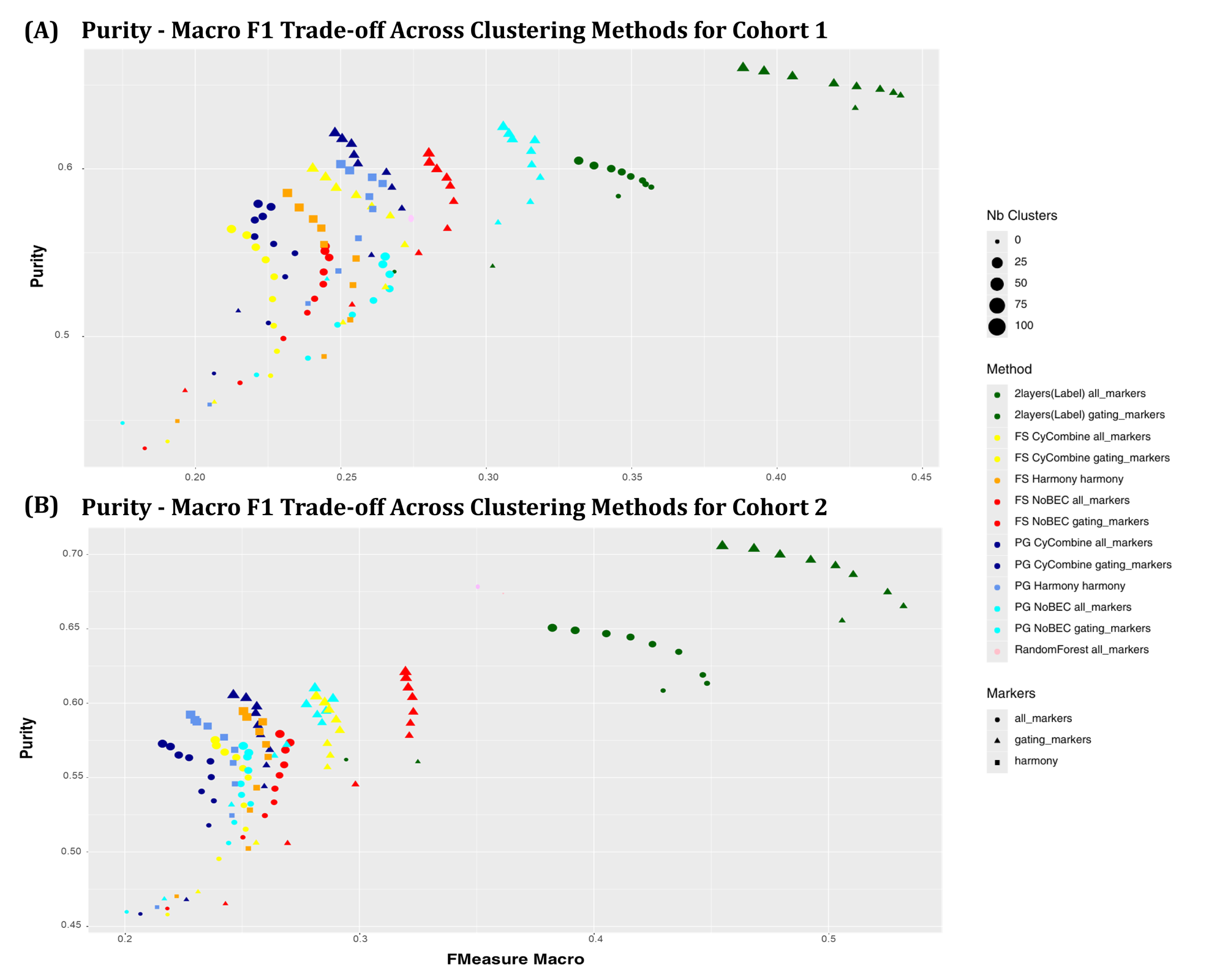
(A) Trade-off between mean purity and Macro-F1 for the resolution of 30 clusters in Cohort 1 (LN); (B) corresponding trade-off in Cohort 2 (SjD).

In both cohorts, FlowSOM and Phenograph variants clustered in the lower-left to middle-left region regardless of BEC condition, achieving high weighted purity through over-fragmentation, or moderate F1 at lower resolutions, but not both. Among the unsupervised methods, PG NoBEC and FS NoBEC with gating markers showed the most competitive profiles, although they remained substantially below AF on both axes. BEC (cyCombine, Harmony) shifted all scatter points leftward relative to NoBEC, confirming its detrimental effect on phenotype recovery.

### 3.4 Rare phenotype recovery: Regulatory T cells (Tregs)

To assess sensitivity for rare functional populations, Treg specific recall was investigated separately across all methods, marker configurations, and cluster resolutions (**Fig. 7A**). In cohort 1, AltraFlowSOM achieved a median Treg recall of 0.29 across cluster resolution 30 using gating markers, compared with 0.25 for PG NoBEC (the next-best method). Beyond the absolute margin, AF demonstrated greater stability across resolutions, whereas other methods showed higher variance, indicating sensitivity to the choice of cluster number. Phenograph variants formed a second performance tier, with moderate but resolution-dependent recall. PG NoBEC exhibited the most competitive profile among the unsupervised methods. FlowSOM-based methods showed near-zero Treg recall across most resolutions, suggesting that the grid-based SOM topology cannot adequately represent rare populations without label guidance. Random Forest also showed near-zero Treg recall across all marker sets despite its overall second-place rank on ARI and Macro-F1, owing to the strong class imbalance. Finally, increasing cluster number did not reliably improve Treg recovery for any method, confirming that rare population detection cannot be addressed by resolution tuning alone and requires a method architecturally designed to anchor low-abundance populations.

**Fig. 7:**
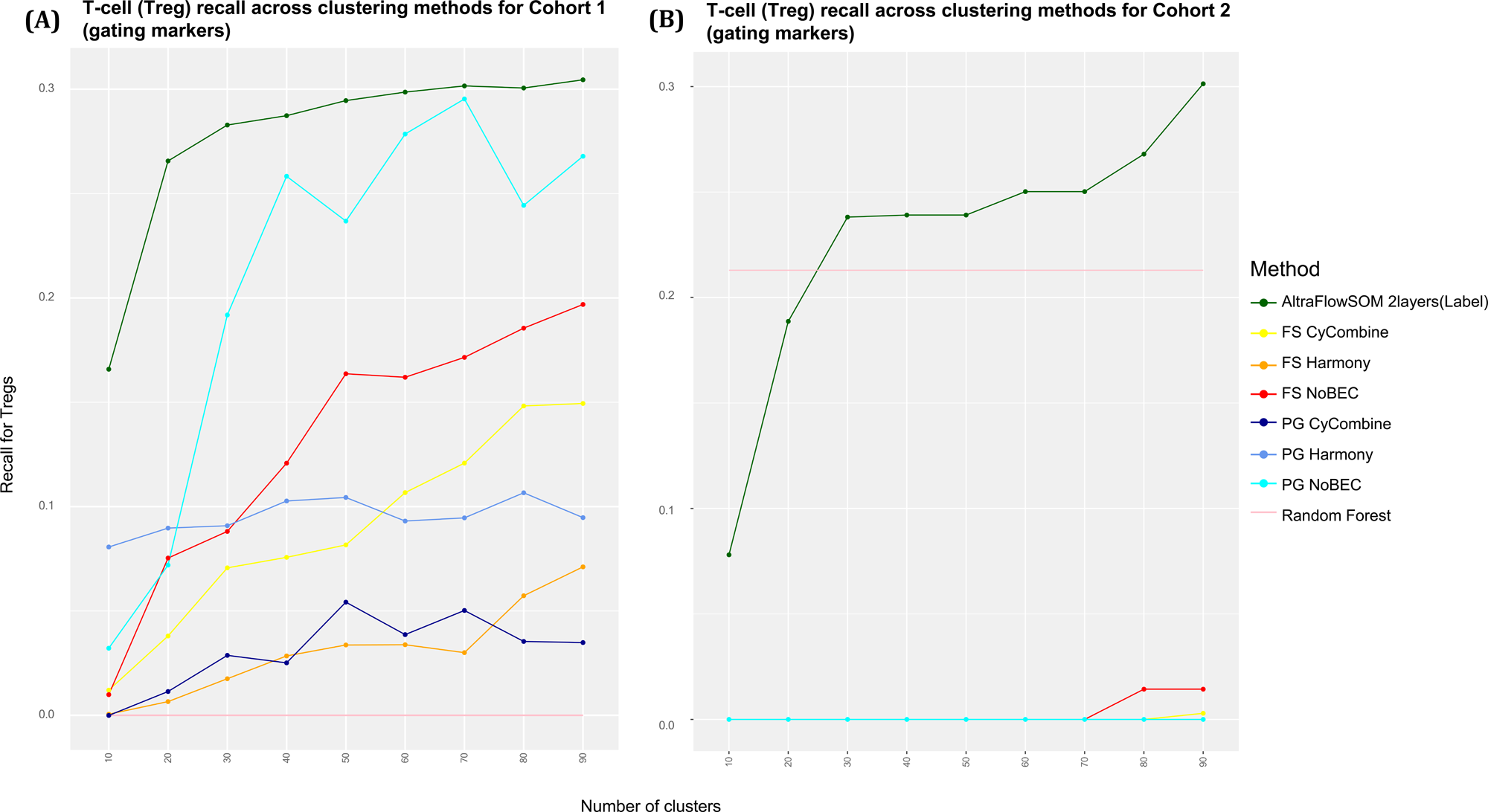
Comparison of regulatory T-cell (Treg) recall across clustering methods using gating-marker configurations. Panel (A) shows Treg recall in Cohort 1, Lupus Nephritis, and panel (B) shows Treg recall in Cohort 2, Sjögren’s disease.

In cohort 2, the recall for regulatory T cells (Tregs) using the gating marker panel revealed pronounced differences between the methods (**Fig. 7B**). The two-layer, AltraFlowSOM substantially outperformed all other unsupervised approaches, with recall increasing from 0.08 at 10 clusters to 0.30 at 90 clusters and surpassing the supervised Random Forest baseline (recall ∼ 0.21) from 30 clusters onward. In contrast, both FlowSOM (FS) and PhenoGraph (PG) pipelines failed to recover Tregs, regardless of the batch-correction strategy applied (CyCombine, Harmony, or no correction), with recall remaining near zero across the entire range of cluster numbers tested; only FS without batch correction showed a marginal increase (∼0.015) at 80-90 clusters. These results indicate that conventional clustering methods, even when combined with batch-effect correction, are unable to resolve the subtle phenotypic distinctions defining Tregs based on gating markers alone, whereas the label-guided hierarchical structure of AltraFlowSOM enables reliable identification of this rare population at sufficient clustering resolution.

## 4. Discussion

AltraFlowSOM was designed to address a fundamental gap in IMC analysis: the inability of existing approaches to simultaneously achieve biological faithfulness, scalability, and rare-population sensitivity within a single reproducible pipeline. Evaluated across two independent IMC cohorts : lupus nephritis kidney biopsies and primary Sjögren’s syndrome salivary gland biopsies, AltraFlowSOM consistently outperformed unsupervised and supervised baselines, demonstrating that semi-supervised SOM training is a viable and generalisable strategy for high-dimensional tissue phenotyping.

IMC uniquely preserves tissue architecture while quantifying dozens of protein markers per cell, positioning it at the intersection of spatial and proteomic information that dissociative approaches (Flow Cytometry, Mass Cytometry (CyTOF), Single-cell RNA Sequencing (scRNA-seq) and CITE-seq) cannot replicate. Cell phenotyping is a prerequisite for any downstream spatial analysis: if the same biological cell type is labelled inconsistently across samples, apparent microenvironmental differences reflect analytical or technical noise rather than true biology. AltraFlowSOM’s higher ARI and Macro-F1 across both cohorts - combined with its unique achievement of simultaneously high recall and purity - confirm that its clusters are internally homogeneous and correctly matched to reference cell populations, without the excessive fragmentation seen in FlowSOM and PhenoGraph variants. The consistency of performance across kidney and salivary gland tissues, which differ substantially in cellular architecture, provides initial evidence that label-guided SOM training generalises across tissue contexts.

The batch-effect correction results require cautious interpretation. In our benchmarking, Harmony and cyCombine did not lead to an overall improvement in clustering performance compared with the uncorrected baselines; instead, they were associated with reduced agreement with the manual reference annotations, as reflected by lower ARI and Macro-F1 scores. Although purity-based measures may remain stable or even increase after correction, this does not necessarily indicate better biological recovery, particularly because weighted purity can be dominated by abundant populations and because purity does not capture recall. Therefore, the observed reduction in ARI and Macro-F1 suggests that batch-effect correction was not beneficial in this setting. One possible explanation is that technical variation was not fully removed, while some biologically informative marker variation may also have been partially attenuated during correction. These findings support the use of multiple complementary metrics, rather than purity alone, when evaluating integration and batch-correction strategies against labelled reference annotations.

The rare phenotype results of AltraFlowSOM are particularly significant. Its median Treg recall of 0.29 exceeded the next-best method and was substantially more consistent across cluster numbers, whereas Random Forest, despite ranking second overall on ARI and Macro-F1 showed near-zero Treg recall, exposing a fundamental limitation of supervised classifiers under class imbalance: when a population such as regulatory T cells constitutes a small minority of cells, discriminative models trained to minimise global loss systematically deprioritise it in favour of abundant classes. The semi-supervised design of AltraFlowSOM directly overcomes this constraint by anchoring SOM topology through partial expert labels while preserving identification - for rare populations, an approach analogous to the transductive learning (Schneider and Xhafa, 2022) paradigm, in which labelled and unlabelled observations jointly shape the decision boundary rather than treating supervision as a post-hoc classification step. More broadly, this result generalises beyond Tregs: any rare population whose clinical significance is disproportionate to its numerical frequency in tissue sections, stands to benefit from methods that embed biological priors into the representation itself rather than applying them after the fact.

A key strength of AltraFlowSOM, although not fully exploited in the present study, is its extensibility beyond the dual-layer (expression + labels) configuration evaluated here (Penn 2005, Kohonen 1995). The SuperSOM framework natively accommodates *L* heterogeneous data layers simultaneously, each with its own distance metric and layer weight w_l_. This modularity makes AltraFlowSOM directly applicable to several emerging experimental designs that represent concrete directions for future work, (a) Transcriptomic and proteomic synergy. Incorporating matched transcriptomic data (e.g. CITE-seq RNA layer) as a third input layer would allow the SOM to resolve activation states that are indistinguishable at the protein level alone, without requiring separate single-cell RNAseq clustering. (b) Spatial and temporal priors: spatial information, could be incorporated as an auxiliary SuperSOM layer alongside marker expression and labels, allowing the physical co-localisation to directly shape map topology and yield niche aware cluster boundaries. A key challenge is that spatial information is not directly comparable across ROIs and would need spatial context features rather than absolute values such as distance to defined structure, local cell density or proximity to a specific cell population and (c) Multi-site harmonization. Encoding the acquisition batch or clinical center as a categorical auxiliary layer would enable batch-aware topology learning without the post-hoc correction step shown here to be detrimental. Realizing these extensions requires datasets with matched multi-modal layers and principled strategies for setting inter-layer weights when the informativeness of each layer varies substantially, and tractable next steps that can be built directly on the already implemented SuperSOM architecture.

This study had several limitations. First, the two-cohort design establishes cross-tissue generalizability within IMC, but true cross-platform validation requires extension to CODEX and MIBI, which share IMC’s conceptual architecture but differ in their signal characteristics and dynamic range. Second, the arcsinh cofactor of 0.1 and gating-marker selection strategy were optimized for these cohorts and may require platform-specific recalibration for panels with different compositions or acquisition settings. Third, manual gating - used here as ground truth - is subject to inter-annotator variability, and AltraFlowSOM’s robustness to label noise has not been formally assessed. Finally, the current evaluation focused on two autoimmune tissue contexts; extension to oncological settings, where tumour heterogeneity and immune infiltration patterns differ substantially, will be necessary to establish broader applicability.

AltraFlowSOM operates on the cell-by-marker intensity matrix produced after segmentation and feature extraction, independently of the acquisition modality, making it directly applicable to any multiplexed imaging platform. At a conceptual level, it resolves a tension that the field has long faced: manual gating is interpretable but not scalable; unsupervised clustering is scalable but not always biologically aligned; supervised classification imposes a closed-world assumption that forecloses discovery. By embedding partial expert annotations directly into SOM topology during training, AltraFlowSOM solves this trilemma without sacrificing any of its three desiderata. The present work establishes semi-supervised SOM training with partial expert annotations as a viable and superior strategy for IMC phenotyping, and provides a benchmarking foundation for future spatial multi-omics integration studies.

## Data Availability

The IMC datasets (cell-by-marker intensity matrices, segmentation masks, and expert manual gating labels) for the lupus nephritis cohort (Cohort 1) and the Sjögren’s syndrome cohort (Cohort 2) analyzed in this study are available from the corresponding authors upon request, subject to applicable institutional and ethical restrictions.

## Code Availability

AltraFlowSOM was developed using the Kohonen package (v3.0.12; Wehrens & Kruisselbrink, 2018; CRAN: https://cran.r-project.org/package=kohonen). The AltraFlowSOM function, along with a sample dataset for testing will be publicly available in the GitHub repository (https://github.com/akhiya96/AltraFlowSOM).

## Acknowledgements

The authors thank the SPATIOMICS platform Patrice Hémon and Yuna Delarue for image acquisition, and Marion Le Rochais for image segmentation and all patients who participated in this study. The authors are grateful to Julien Nourikyan, Saloni Patel, and Vivekanand Asokachandran for their scientific discussions, and to Rinu Roy for unwavering support. AI-assisted tools such as PaperPal (Cactus Communications) and Claude AI (Anthropic) for language editing were used during the manuscript preparation. All scientific content and conclusions are the sole responsibility of the authors. The authors acknowledge the cytometry flow core facility Hyperion (Brest, France) for its technical assistance, as well as the European FEDER grant program Progos RU 000950 and members of the Scientific Interest Group (GIS) Biogenouest.

## Funding Info

This study was conducted with the support of the European Union, French State, Région Bretagne, Conseil Départemental du Finistère and Brest Métropole within the framework of the CPER B2S. This project has received funding from the European Union’s Horizon 2020 Research and Innovation Programme under Grant Agreement No 731944.’ Content of this publication reflects the author’s view and the Commission is not responsible for any use that may be made of the information it contains.

## Author Contributions

**A.A.R.:** Conceptualization, Methodology, Software, Formal analysis, Data curation, Visualization, Writing original draft, Writing review and editing. **E.B.:** Resources, Investigation, Writing : review and editing. **C.L.D.:** Resources, Investigation, Formal analysis, Writing: review and editing. **P.H.:** Resources, Writing : review and editing. **P.E.J.:** Conceptualisation, Methodology, Software, Supervision, Writing: review and editing. **S.H.:** Conceptualisation, Project administration, Resources, Supervision, Writing : review and editing.

## Ethics approval and patient consent

The study was performed in accordance with the Declaration of Helsinki and was approved by the relevant ethical committees. The Diagnostic Suspicion of Primitive Syndrome Sjogren’s (DIApSS) study is registered at [https://clinicaltrials.gov/study/NCT03681964 | ClinicalTrials.gov] (identifier: NCT03681964). The Systemic Lupus Erythematosus Within the Framework of Multidisciplinary Consultation (CO LUPUS) study is registered at [https://clinicaltrials.gov/study/NCT04320680] (identifier: NCT04320680).

